# Distinct neuronal processes in the ventromedial prefrontal cortex mediate changes in attention load and nicotine pro-cognitive effects

**DOI:** 10.1101/2024.10.15.618452

**Authors:** Caroline Vouillac-Mendoza, Nathalie Biendon, Sandra Dovero, Karine Guillem

## Abstract

The prefrontal cortex (PFC) plays a key role in attention. In particular, neuronal activity in the ventromedial PFC (vmPFC) has been implicated in the preparatory attentional period that immediately precedes cue presentation. However, whether vmPFC neuronal activity during this preparatory period is also sensitive to changes in task demand and to the pro-cognitive effects of nicotine remained to be investigated. Here, we used *in vivo* electrophysiology to record vmPFC neuronal activity during two distinct manipulations: a task manipulation that increased task demand by reducing the cue stimulus duration (from 1s to 0.5s), and a pharmacological manipulation by administrating an acute nicotine injection (10 µg/inj, i.v.) before the session. We found that increasing task demand decreased attentional performances and vmPFC precue neuronal activity, but had no effect on gamma oscillations. In contrast, nicotine injection increased attention and gamma oscillations, but almost abolished vmPFC phasic precue responses. Together, these findings indicate the existence of two distinct neuronal mechanisms operating at different timescales and suggests that allocation of attention could be achieved through multiple complementary neuronal mechanisms within the vmPFC.

## INTRODUCTION

The prefrontal cortex (PFC) plays a critical role in attentional processes in humans and animals (Christakou et al., 2004; Kim et al., 2016; Miller and Cohen, 2001; Totah et al., 2009), and dysfunctions of the PFC have been implicated in several disorders, including attentional deficit hyperactivity disorder (ADHD), Alzheimer’s disease or schizophrenia (Chudasama and Robbins, 2006; Dalley et al., 2004; Halperin and Schulz, 2006). In rodents, PFC-dependent cognitive functions have been extensively studied using the 5-choice serial reaction time task (5-CSRTT), a well-established test setup that taxes various aspects of attentional control over performance (Bari et al., 2008; Robbins, 2002).

Previous electrophysiologic single-unit recordings in both primates and rodents have implicated mPFC activity during attention tasks (Gill et al., 2000; Johnston et al., 2007; Niki and Watanabe, 1979; Terra et al., 2020). Specifically, these studies have demonstrated that attention-related mPFC neuronal activity typically occurs in anticipation of oncoming task-relevant visual cues on a time scale of seconds (Donnelly et al., 2015; Kim et al., 2016; Totah et al., 2013; Totah et al., 2009). Moreover, theses precue responses appear to be related to the degree of preparatory attention and subsequent response (Donnelly et al., 2015; Kim et al., 2016; Totah et al., 2009). Though they all participate in attention, evidence suggest that distinct mPFC subregions contribute to separate aspects of attentional processing and are required at distinct time windows during preparatory attentional states (de Kloet et al., 2021; Hardung et al., 2017; Luchicchi et al., 2016; Terra et al., 2020; Totah et al., 2013; Totah et al., 2009). In particular, the ventromedial PFC (vmPFC) neuronal activity plays an important role during the seconds that immediately precede cue presentation, as transient inhibition of vmPFC pyramidal neurons during that precue preparatory period impairs 5-CSRTT performances in rats (Luchicchi et al., 2016).

Several factors or task manipulations have been shown to alter attentional behavior. For instance, increasing task demand by reducing the stimulus duration during a 5-CSRTT session decreased attention, while in contrast nicotine administration before the session increased it (Amitai and Markou, 2009; Hahn et al., 2002; Mirza and Stolerman, 1998; Robbins, 2002). One open question, however, is whether vmPFC neuronal activity is sensitive to changes in task demand and to the pro-cognitive effects of nicotine. Here, we used *in vivo* electrophysiological recordings in rats trained to perform an attentional 5-CSRTT task to directly assess this question. We used two distinct manipulations with opposite effect on attentional performance: one that reduced attention by reducing the stimulus duration from 1s to 0.5s (i.e., SD1s or SD0.5s), and the other that increased attention by administrating an acute nicotine injection (10 µg/inj, i.v.) before the session. We then recorded and compared *in vivo* vmPFC neuronal activity during these two manipulations.

## RESULTS

### Attentional performances vary with attentional load

We first determined animals’ attentional capacities using the five-choice serial reaction time task (5-CSRTT). Briefly, animals (*n* = 8) were trained to detect and respond to a brief light stimulus randomly presented in one of five nose poke holes to receive a sucrose reward (0.1 ml of 10% sucrose), as previously described (Vouillac-Mendoza et al., 2024) (see Materials and Methods for details; Fig. 1a, b). During the training schedule, the stimulus duration (SD) progressively decreased across sessions until they reached stable performance with a final SD 1s (from 30s to 1s, reached after 36 ± 2 training sessions). To further test attention, attentional load was then increased using a SD variable procedure, in which stimulus durations decreased from 1 to 0.5 seconds within the same session. Animals were trained under this SD variable procedure for 10 more days until they reached stable performance. As expected, reducing the SD duration from 1s to 0.5s impaired the attentional performances of the rats (*n* = 8) and resulted in a significant decrease in accuracy (from 78.7 ± 2.0% to 69.1 ± 3.4% for SD1s and SD0.5s respectively; *F*_(1,7)_ = 11.23; *p* < 0.01; Fig. 1c), and a small but not significant increase in the percentage of omission (*F*_(1,7)_ = 3.91; *NS*; Fig. 1d). The decrease in accuracy during SD0.5s was mainly caused by an increase in incorrect responses (*F*_(1,7)_ = 12.81; *p* < 0.01; Fig. 1f), rather than a decrease in correct responses (*F*_(1,7)_ = 1.14; *NS*; Fig. 1e). Furthermore, there was no effect of SD duration on any other all behavioral measures, such as number of premature responses (*F*_(1,7)_ = 1.6; NS), correct response latency (*F*_(1,7)_ = 2.4; NS), incorrect response latency (*F*_(1,7)_ = 3.1; NS), or latency to collect sucrose reward (*F*_(1,7)_ = 0.4; NS; Table 1), suggesting that decreased accuracy reflected impairments in attention processes rather than motor or motivational deficits.

**Figure 1.**
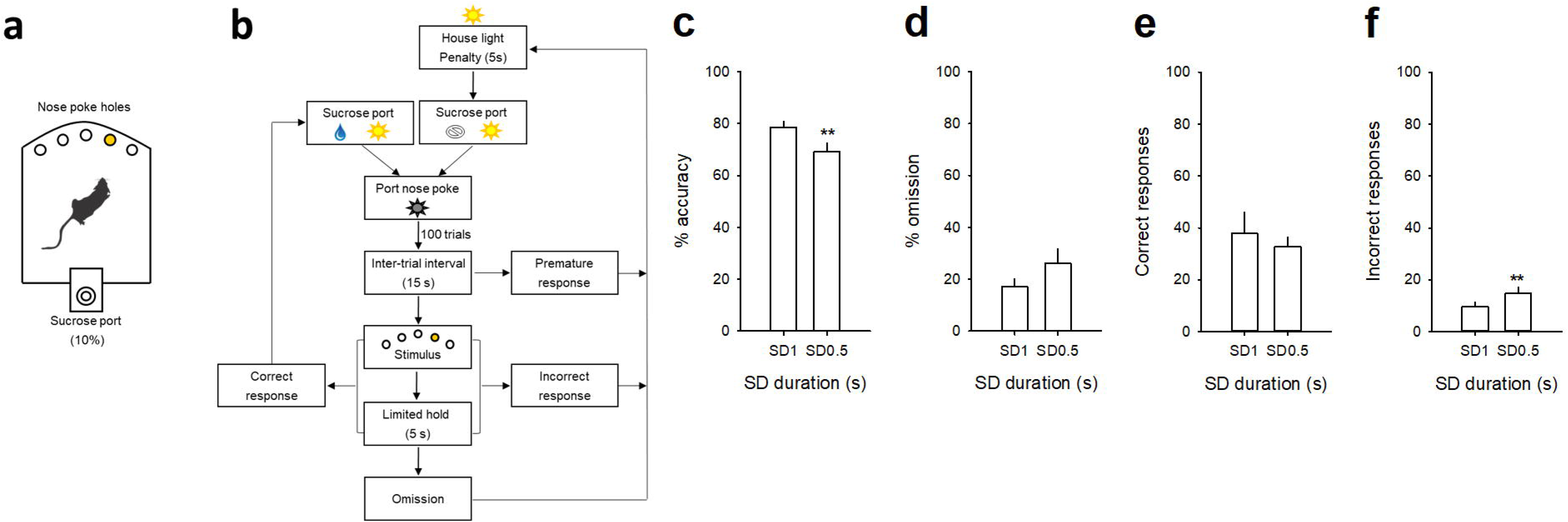
Attentional performances in 5-CSRTT during a stimulus duration (SD) variable procedure. (**a**) Schematic of the 5-CSRTT apparatus and (**b**) diagram showing the sequence of events during a 5-CSRTT training session. Rats initiate trials by responding into the sucrose port; after an intertrial interval delay, a brief stimulus is presented in one of the five holes. Subjects must respond to that hole (correct response) within a certain time (limited hold) to receive a sucrose reward (0.1 ml of 10% sucrose). If they respond to the wrong hole (incorrect response), respond before the stimulus is presented (premature response), or fail to respond (omission), they are punished with a penalty period during which the house light was turned on. After that, the sucrose port is switched on and rats have to respond into it to initiate the next trial. (**c, d**) Mean (±SEM) percentage of accuracy (**c**), omission (**d**) and (**e, f**) mean (±SEM) number of correct (**e**) and incorrect (**f**) responses as a function of the light cue stimulus duration, respectively SD1s and SD 0.5s. \*\**p* < 0.01, different from SD1s.

**Table 1:** 5-CSRTT performances at varying stimulus durations

|  | Stimulus durations (s) |  |
| --- | --- | --- |
|  | SD 1s | SD 0.5s |
| Premature | $7.8 \pm 2.5$ | $5.5 \pm 1.4$ |
| Correct latency (s) | $1.1 \pm 0.0$ | $1.0 \pm 0.1$ |
| Incorrect latency (s) | $1.4 \pm 0.1$ | $1.2 \pm 0.1$ |
| Reward latency (s) | $1.5 \pm 0.1$ | $1.6 \pm 0.2$ |

### vmPFC neuronal phasic activity is associated with changes in attentional performances

We next recorded i*n vivo* vmPFC neuronal activity during this SD variable procedure in the same animals. Based on previous reports (Kim et al., 2016; Totah et al., 2013; Totah et al., 2009), we focused our examination on the responses of vmPFC pyramidal neurons recorded during the precue period (−2 to 0 s before cue onset), when attention is allocated. vmPFC neurons whose firing activity changed during the precue period were identified for each SD duration, and neuronal responses were divided into three groups depending on whether a correct response, an incorrect response, or an omission of response followed. Precue-related phasic responses were observed on each trial and consisted of mixed changes in activity during the precue period with half of the neurons showing an increase and half of the neurons showing a decrease in firing activity on every trials type and SD durations. However, the proportion of phasic precue responses were different depending on trial type and SD duration (Fig. 2). Table 2 lists the number and percentage of precue responsive neurons on each trial type (correct, incorrect and omission trials) and for each SD duration (SD1s or SD 0.5s). Notably for the SD1s with high attentional performances, the proportion of precue responsive neurons was higher on correct trials than on incorrect (57% vs 30%, Z = 4.24; p < 0.001; Fig. 2b, c and Table 2) or omissions trials (57% vs 22%, Z = 5.61; p < 0.001; Fig. 2b, d and Table 2). For the shorter SD0.5s with low attentional performances in contrast, the proportion of precue responsive neurons was similar on correct and incorrect trials (48% vs 40%, Z = 1.29; NS Fig. 2 b, c and Table 2), while still higher than on omission trials (48% vs 23%, Z = 4.28; p < 0.001; Fig. 2 b, d and Table 2). The weakened difference between the proportion of correct and incorrect precue responsive neurons observed when the SD decrease to 0.5s was mainly caused by an increased responding during incorrect trials (30% vs 40%, Z = 1.60; p < 0.05) rather than a decreased responding during correct trials (57% vs 48%, Z = 1.40; p = 0.08).

**Figure 2.**
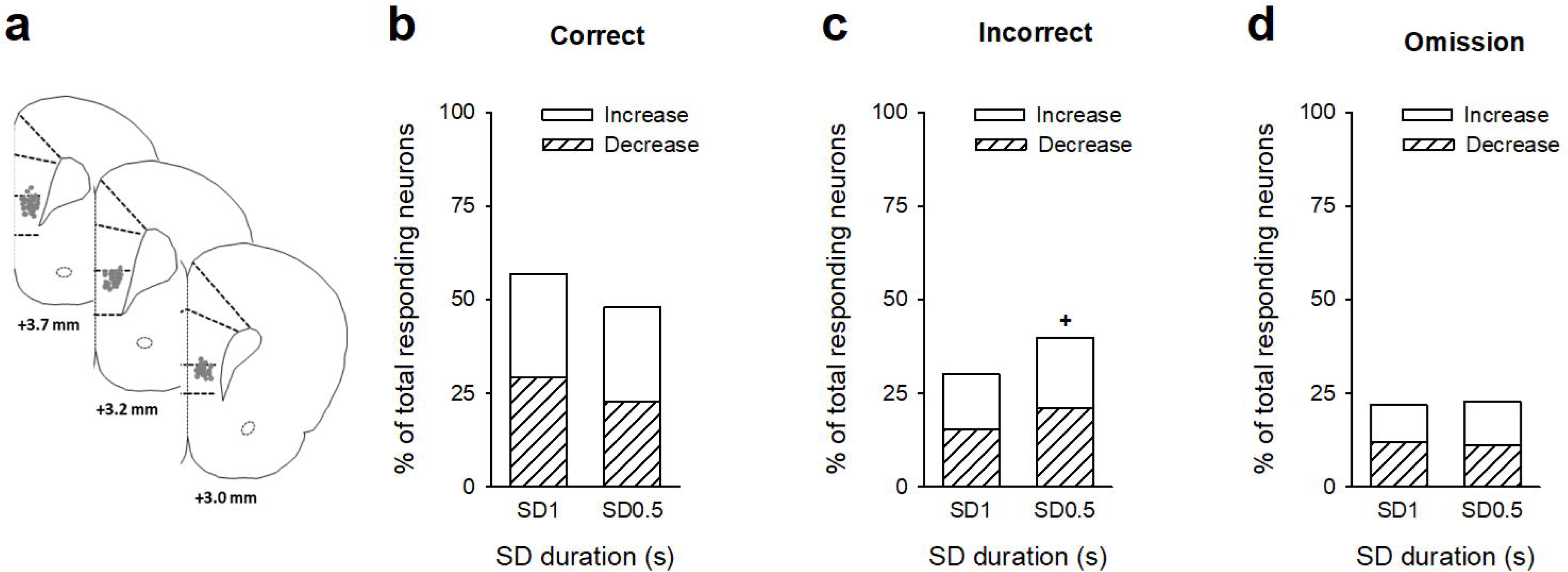
vmPFC precue phasic responses during the SD variable procedure. (**a**) Schematic of the location of individual wire tips within the vmPFC. The numbers indicate millimeters anterior to bregma. (**b, c, d**) Percentage of total neurons responding before the cue on correct (**b**), incorrect (**c**), and omission trials (**d**) as a function of the SD duration, respectively SD1s and SD 0.5s. Excitatory units are shown as solid bars, and inhibitory units are shown as hashed bars. ^+^*p* < 0.05, different from SD1s.

**Table 2:** Number and percentage of vmPFC neurons responding during the precue period.

|  | Correct trials |  | Incorrect trials |  | Omission trials |  |
| --- | --- | --- | --- | --- | --- | --- |
|  | SD 1s | SD 0.5s | SD 1s | SD 0.5s | SD 1s | SD 0.5s |
| Total | 70 (60%) | 59 (48%) | 37 (30%)*** | 49 (40%)<sup>+</sup> | 27 (22%)*** | 28 (23%)*** |
| Increase | 34 (27%) | 31 (25%) | 18 (15%)** | 23 (19%) | 18 (15%)** | 14 (11%)** |
| Decrease | 36 (29%) | 28 (23%) | 19 (15%)** | 26 (21%) | 19 (15%)** | 14 (11%)** |

\*\*\* $p < 0.001$ and \*\* $p < 0.01$ , significant difference compared to correct trials. <sup>++</sup> $p < 0.01$ and <sup>+++</sup> $p < 0.001$ , significant difference compared to SD 1s.

Similarly, the magnitude of precue responses were graded depending on trial type and SD duration (Fig. 3). For the SD1s, the vmPFC pyramidal subpopulation with increased activity in correct trials displayed lower firing rates in incorrect and omission trials (time X trial type: F(118, 5546) = 1.82; *p* < 0.001; Fig. 3a and trial type: F(2, 94) = 13.52; *p* < 0.001; Correct *vs* Incorrect, *p* < 0.01; Correct *vs* Omission, *p* < 0.001 and Incorrect *vs* Omission, *NS*; Fig. 3c). Inversely, the vmPFC pyramidal subpopulation with decreased activity in correct trials was less suppressed in incorrect and omission trials (time X trial type: F(118, 5782) = 2.82; *p* < 0.001; Fig. 3a and trial type: F(2, 98) = 21.24; *p* < 0.001; Correct *vs* Incorrect, *p* < 0.001; Correct *vs* Omission, *p* < 0.001 and Incorrect *vs* Omission, *NS*; Fig. 3c). The magnitude of precue responses were also graded at the shorter SD0.5s for both the increased (time X trial type: F(118, 5133) = 3.03; *p* < 0.001; Fig. 3b and trial type: F(2, 87) = 5.94; *p* < 0.01; Correct *vs* Incorrect, *p* < 0.05; Correct *vs* Omission, *p* < 0.01 and Incorrect *vs* Omission, *NS*; Fig. 3d) and the decreased vmPFC pyramidal subpopulations (time X trial type: F(118, 4661) = 2.05; *p* < 0.001; Fig. 3b and trial type: F(2, 79) = 6.36; *p* < 0.01; Correct *vs* Incorrect, *p* < 0.05; Correct *vs* Omission, *p* < 0.01 and Incorrect *vs* Omission, *NS*; Fig. 3d), but to a lesser extent than in SD1s for the correct trials (SD X trial type: F(2, 180) = 7.08; *p* < 0.001; SD1 vs SD0.5, *p* < 0.001 and SD X trial type: F(2, 177) = 8.29; *p* < 0.001; SD1 vs SD0.5, *p* < 0.001, for the increased and the decreased subpopulations respectively). Finally, at the population level of all recorded neurons, the neuronal activity during the precue period for all trials combined was significantly higher in SD1s than in SD0.5s (F(1, 122) = 5.16; *p* < 0.05; Fig. 3e). Together, these data indicated that changes in attentional load induced changes in attentional performances and associated vmPFC pyramidal precue neuronal activity: the lower the task demand (i.e., SD1s), the higher the performance and the higher vmPFC precue firing activity.

**Figure 3.**
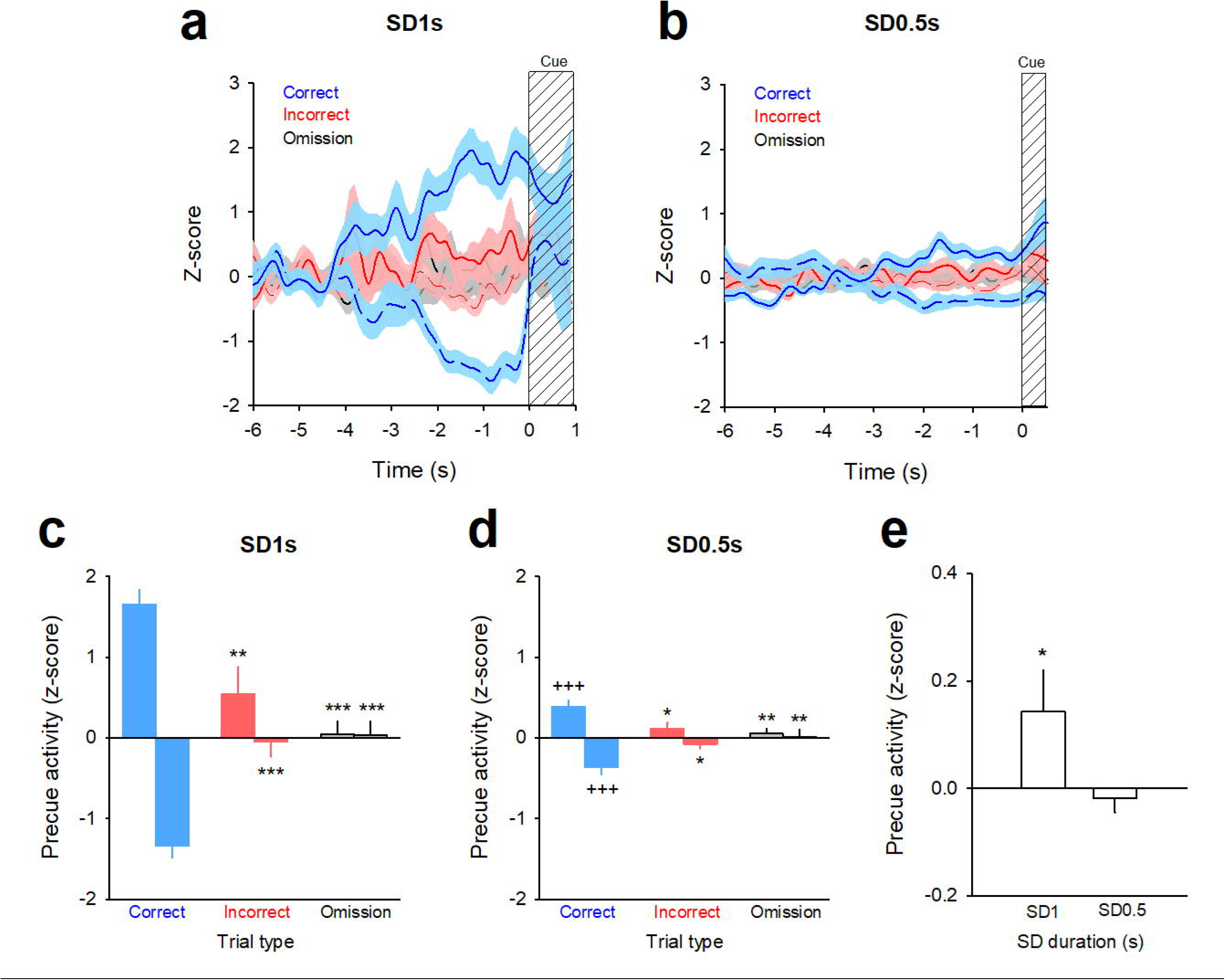
Firing activity of vmPFC precue responsive neurons before the SD1s and SD0.5s cue presentation. (**a**, **c**) Mean (±SEM) normalized firing rate (z-scores smoothed with a Gaussian filter) of vmPFC precue responses (excitatory, solid line; inhibitory, dashed line) on correct trials (*blue*) and the activity of those same units on incorrect (*red*) and omission (*grey*) trials before the SD1s (**a**) and SD0.5s (**c**) cue presentation. The lines represent the mean activity and the shaded areas represent the SEM. (**b**, **d**) Mean magnitude (±SEM) of precue activity (z-score) on correct (*blue*), incorrect (*red*) and omission (*grey*) trials before the SD1s (**b**) and SD0.5s (**d**) cue presentation. \**p* < 0.05 and \*\**p* < 0.01, different from correct trials. ^+++^*p* < 0.001, different from SD1s. (**e**) Mean magnitude (±SEM) of precue activity (z-score) of all recorded neurons for all trials combined during SD1s and SD0.5s. \**p* < 0.05, different from SD1s.

### vmPFC opto-inhibition differently affects performances depending on attentional load

Based on these observations, we hypothesized that the contribution of vmPFC precue neuronal activity will vary depending on the attentional load and that inhibition of vmPFC neurons should differently impact attention depending on the task difficulty. To test this hypothesis, an adeno-associated virus (AAV) targeting the pyramidal neurons and encoding the archaerhodopsin (CaMKII-ArChT3.0-eYFP; *n* = 12) was bilaterally injected into the vmPFC of a separate group of rats with bilateral implantation of chronic optic fibers in the vmPFC (see Material and Methods for details; Fig. 4a). As a control, we used a virus expressing YFP only (*n* = 8). The efficacy of this in vivo viral strategy was demonstrated by confocal analysis showing that eYFP preferentially colocalized with CaMKII (Fig. 4b) and indicated efficient transduction in pyramidal neurons (selectivity index > 85%). Efficiency of optical stimulation in inhibiting vmPFC activity was demonstrated by cFos immunochemistry showing that light application selectively reduced cFos expression in ArChT but not in YFP rats (F(1,16) = 10.76; *p* < 0.01; Fig. 4c, d). At the behavioral level, opto-inhibition of vmPFC pyramidal neurons 2 s prior to cue presentation differentially alters attentional performance depending on the task demand. Consistent with previous studies (Luchicchi et al., 2016), at the long SD1s associated with high attentional performances, vmPFC opto-inhibition impaired attention as it decreased the accuracy (light X virus interaction: F(1,18) = 7.05; *p* < 0.05; ArChT-No light *vs* ArChT-Light, *p* < 0.001; Fig. 4f) but not the percentage of omission (light X virus interaction: F(1,18) = 0.75; *NS*; Fig. 4g). This effect was due to a decrease in the number of correct responses (light X virus interaction: F(1,18) = 6.25; *p* < 0.05; ArChT-No light *vs* ArChT-Light, *p* < 0.01; Table 3) and a concomitant increase in the number of incorrect responses (light X virus interaction: F(1,18) = 4.90; *p* < 0.05; ArChT-No light *vs* ArChT-Light, *p* < 0.001; Table 3). In contrast, at the shorter SD0.5s associated with lower attentional performance, vmPFC opto-inhibition decreased the percentage of omission (light X virus interaction: F(1,18) = 7.05; *p* < 0.05; ArChT-No light *vs* ArChT-Light, *p* < 0.001; Fig. 4i) without affecting the accuracy (light X virus interaction: F(1,18) = 0.57; *NS*; Fig. 4h). This decrease in omission was caused by a greater engagement of the rats in the task as they performed more correct (light X virus interaction: F(1,18) = 4.30; *p* < 0.05; ArChT-No light *vs* ArChT-Light, *p* < 0.01; Table 3) and incorrect responses (light X virus interaction: F(1,18) = 15.68; *p* < 0.001; ArChT-No light *vs* ArChT-Light, *p* < 0.001; Table 3), suggesting an increase in general arousal. Optogenetic inhibition of vmPFC pyramidal neurons did not affect le number of premature responses at any SD duration (light X virus interaction: F(1,18) = 0.23; and F(1,18) = 1.65; *NS*; for SD1s and SD0.5s, respectively; Table 3). Light application also reduced correct (light: F(1,18) = 21.9; and F(1,18) = 42.0; *p* < 0.001; for SD1s and SD0.5s, respectively; Table 3) and incorrect latencies (light: F(1,18) = 21.9; and F(1,18) = 24.5; *p* < 0.001; for SD1s and SD0.5s, respectively; Table 3) at each SD duration, suggesting that rats may detect the optical stimulation which may act as a cue that accelerates response latency. Importantly however, this decrease in response latencies was observed to a similar extend in both YFP and ArChT rats (correct latency : light X virus interaction: F(1,18) = 0.17; and F(1,18) = 1.27; *NS*; for SD1s and SD0.5s, respectively; and incorrect latency : light X virus interaction: F(1,18) = 2.23; and F(1,18) = 0.68; *NS*; for SD1s and SD0.5s, respectively; Table 3) and could thus not explain *per se* the changes in attentional performances.

**Figure 4.**
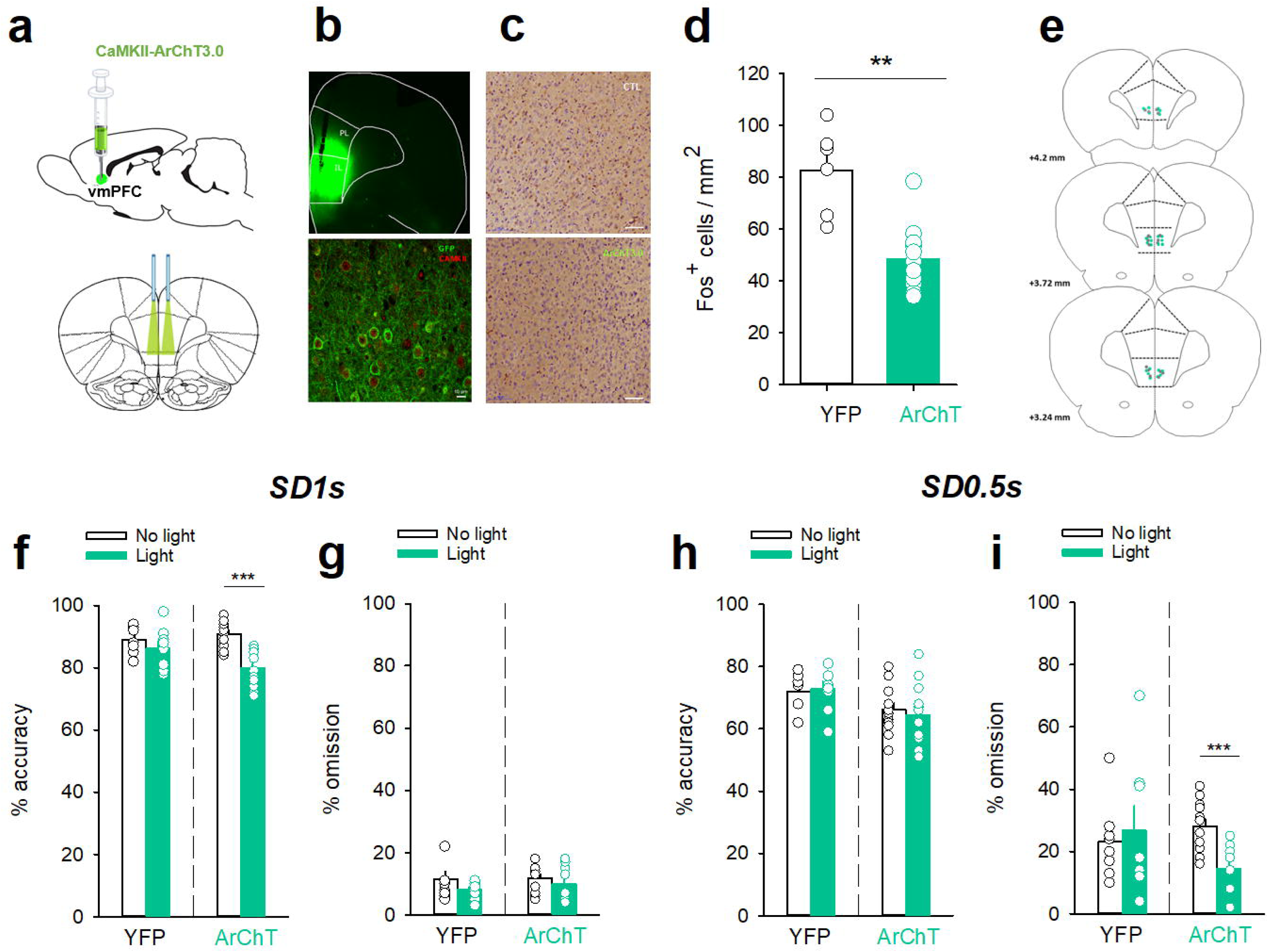
vmPFC optogenetic inhibition before the SD1s and SD0.5s cue presentation differentially altered attention. (**a**) Graphic representations of the injections made in the vmPFC (*top*) and of the bilateral illumination of the vmPFC via patch cables (*bottom*). (**b**) Representative images of coronal sections showing the virus expression and fibers implants in the vmPFC (*top*) and immunostaining of eYFP (green) expression in vmPFC pyramidal neurons (red, CAMKII) (*bottom*). (**c**) Representative photomicrographs of cFos labelling in the vmPFC after the opto-inhibition of vmPFC neurons in rats injected with YFP (*top*) or with ArChT (*bottom*). (**d**) Mean (±SEM) density of Fos-positive cells (cFos+ cells/mm2) in the vmPFC after the opto-inhibition of the vmPFC in rats injected with YFP (*white bars*) or with ArChT (*green bars*). \*\**p* < 0.01, different from YFP. (**e**) Schematic of the location of individual optic fiber tips within the vmPFC. The numbers indicate millimeters anterior to bregma. (**f** - **i**) Mean (±SEM) percentage of accuracy (**f**, **h**) and omission (**g**, **i**) in animals injected either with YFP (n = 8) or ArChT3.0 (n = 12) under light or no-light conditions during the vmPFC inhibition 2 s before the SD1s (**f**, **g**) or SD0.5s (**h**, **i**) cue presentation. \*\*\**p* < 0.0 1, different from No Light.

**Table 3:** Effects of optogenetic inhibition of vmPFC pyramidal neurons on 5-CSRTT performances at varying stimulus durations

|  | SD 1s |  |  |  | SD 0.5s |  |  |  |
| --- | --- | --- | --- | --- | --- | --- | --- | --- |
|  | YFP |  | ArChT |  | YFP |  | ArChT |  |
|  | No light | Light | No light | Light | No light | Light | No light | Light |
| Correct responses | 77.0 ± 2.5 | 76.8 ± 2.4 | 76.2 ± 1.9 | 67.3 ± 2.4** | 44.8 ± 3.2 | 45.0 ± 5.4 | 40.6 ± 2.3 | 50.2 ± 3.6** |
| Incorrect responses | 9.2 ± 1.4 | 11.4 ± 2.1 | 7.6 ± 1.0 | 17.0 ± 1.5*** | 15.3 ± 1.7 | 14.4 ± 2.0 | 20.5 ± 1.2 | 27.1 ± 2.0*** |
| Premature | 4.3 ± 1.5 | 5.8 ± 1.7 | 5.1 ± 0.9 | 6.5 ± 1.3 | 17.9 ± 2.7 | 14.4 ± 1.9 | 14.6 ± 2.2 | 9.8 ± 1.7 |
| Correct latency (s)<sup>+++</sup> | 1.1 ± 0.1 | 1.0 ± 0.0 | 1.0 ± 0.0 | 0.9 ± 0.1 | 1.2 ± 0.0 | 1.1 ± 0.1 | 0.9 ± 0.0 | 0.8 ± 0.0 |
| Incorrect latency (s)<sup>+++</sup> | 1.5 ± 0.1 | 1.2 ± 0.1 | 1.5 ± 0.1 | 0.7 ± 0.1 | 1.3 ± 0.1 | 1.0 ± 0.1 | 1.3 ± 0.1 | 0.8 ± 0.0 |
| Reward latency (s) | 1.3 ± 0.1 | 1.3 ± 0.1 | 1.2 ± 0.1 | 1.2 ± 0.0 | 1.4 ± 0.1 | 1.5 ± 0.1 | 1.3 ± 0.1 | 1.4 ± 0.1 |

Data are presented as mean ± SEM. \* $p < 0.05$ , \*\* $p < 0.01$ , and \*\*\* $p < 0.001$ significant difference compared to No light. +++ $p < 0.001$ , global significant effect of Light.

### Acute nicotine administration increases attention and potentiates vmPFC neuronal tonic and gamma activities

Because nicotine has well known pro-cognitive properties (Levin et al., 2006; Mirza and Stolerman, 1998), we next assessed the effect of acute non-contingent administration of nicotine (10 µg/inj, i.v., 1 min before the session) on attentional performances and associated vmPFC neuronal activity (Fig. 5). As expected, nicotine increased the accuracy at both SD durations (nicotine effect: F(1, 6) = 9.90; *p* < 0.05 and nicotine X SD interaction: F(1, 6) = 0.05; *NS*; Fig. 5a), without affecting the percentage of omission (nicotine effect: F(1, 6) = 2.08; *NS* and nicotine X SD interaction: F(1, 6) = 3.40; *NS*; Fig. 5b). Nicotine had no significant effect on any other all behavioral measures (Table 4), such as number of premature responses (F(1,6) = 0.31; *NS*), correct response latency (F(1,6) = 0.03; *NS*), incorrect response latency (F(1,6) = 0.15; *NS*), or latency to collect sucrose reward (F(1,6) = 0.22; *NS*). Surprisingly however, this nicotine treatment almost abolished vmPFC phasic precue responses (Fig. 6a-c and Table 5) and had no effect on the precue neuronal activity of all recorded neurons (nicotine effect: F(1,60) = 0.09; *NS*; nicotine X SD interaction: F(1,60) = 0.01; *NS*; Fig. 6d). Indeed, the percentage of precue responsive neurons strongly decreased at both SD durations for correct (57% vs 18%, Z = 5.29; *p* < 0.001 and 48% vs 17%, Z = 5.19; *p* < 0.001; for SD1s and SD0.5s respectively; Fig. 6a), incorrect (30% vs 16%, Z = 2.57; *p* < 0.01 and 40% vs 10%, Z = 5.43; *p* < 0.001; for SD1s and SD0.5s respectively; Fig. 6b) and omissions trials (22% vs 8%, Z = 3.01; *p* < 0.01 and 23% vs 7%, Z = 3.58; *p* < 0.001; for SD1s and SD0.5s respectively; Fig. 6c). This lost of precue phasic responses seems due to an overall basal tonic increase in firing rate of all vmPFC recorded neurons after nicotine injection (F(1,53) = 20.21; *p* < 0.001; Fig. 6e), an effect that could increase the background firing rate thus limiting the detection of phasic changes in firing and increased in precue neuronal activity (Kravitz and Peoples, 2008; Peoples et al., 2007). In contrast, while increasing the attentional load had no effect on baseline vmPFC gamma power density (SD effect: F(1,6) = 1.58; *NS*; Fig. 7a-c) or the percentage of gamma oscillations (SD effect: F(1,6) = 0.77; *NS*; Fig. 7d), acute nicotine treatment increased vmPFC gamma power density (nicotine effect: F(1,6) = 8.10; *p* < 0.05; nicotine X SD interaction: F(1,6) = 0.16; *NS*; Fig. 7c) and the percentage of gamma oscillations (nicotine effect: F(1,6) = 7.85; *p* < 0.05; nicotine X SD interaction: F(1,6) = 0.09; *NS*; Fig. 7d) at both SD durations.

**Figure 5.**
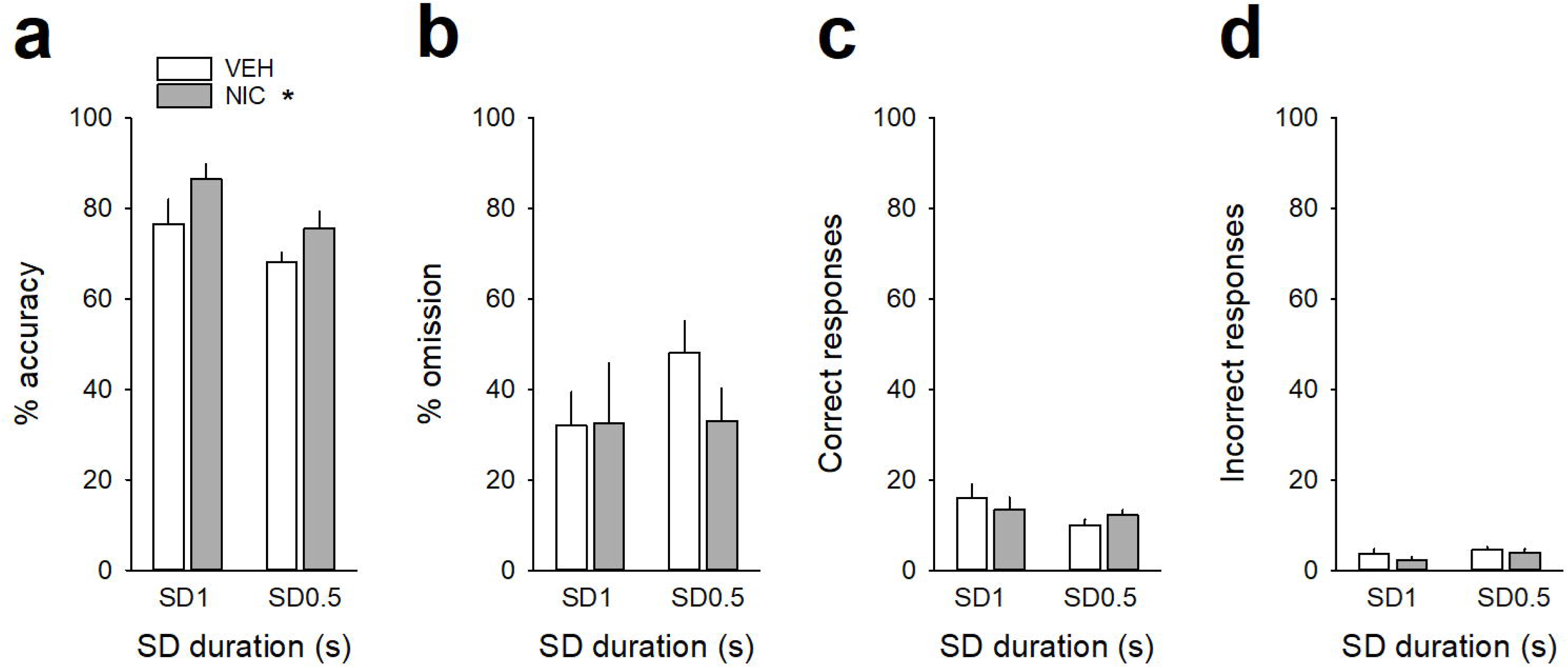
Acute nicotine treatment increases attention. (**a**, **b**) Mean (±SEM) percentage of accuracy (**a**), omission (**b**) and (**c, d**) mean (±SEM) number of correct (**c**) and incorrect (**d**) responses after a saline injection (VEH, *white bars*) and after a nicotine challenge session (NIC, 10 µg/injection, i.v. 1 min before the session, *grey bars*) during the SD1s and SD0.5s cue presentation. \**p* < 0.05, different from VEH.

**Figure 6.**
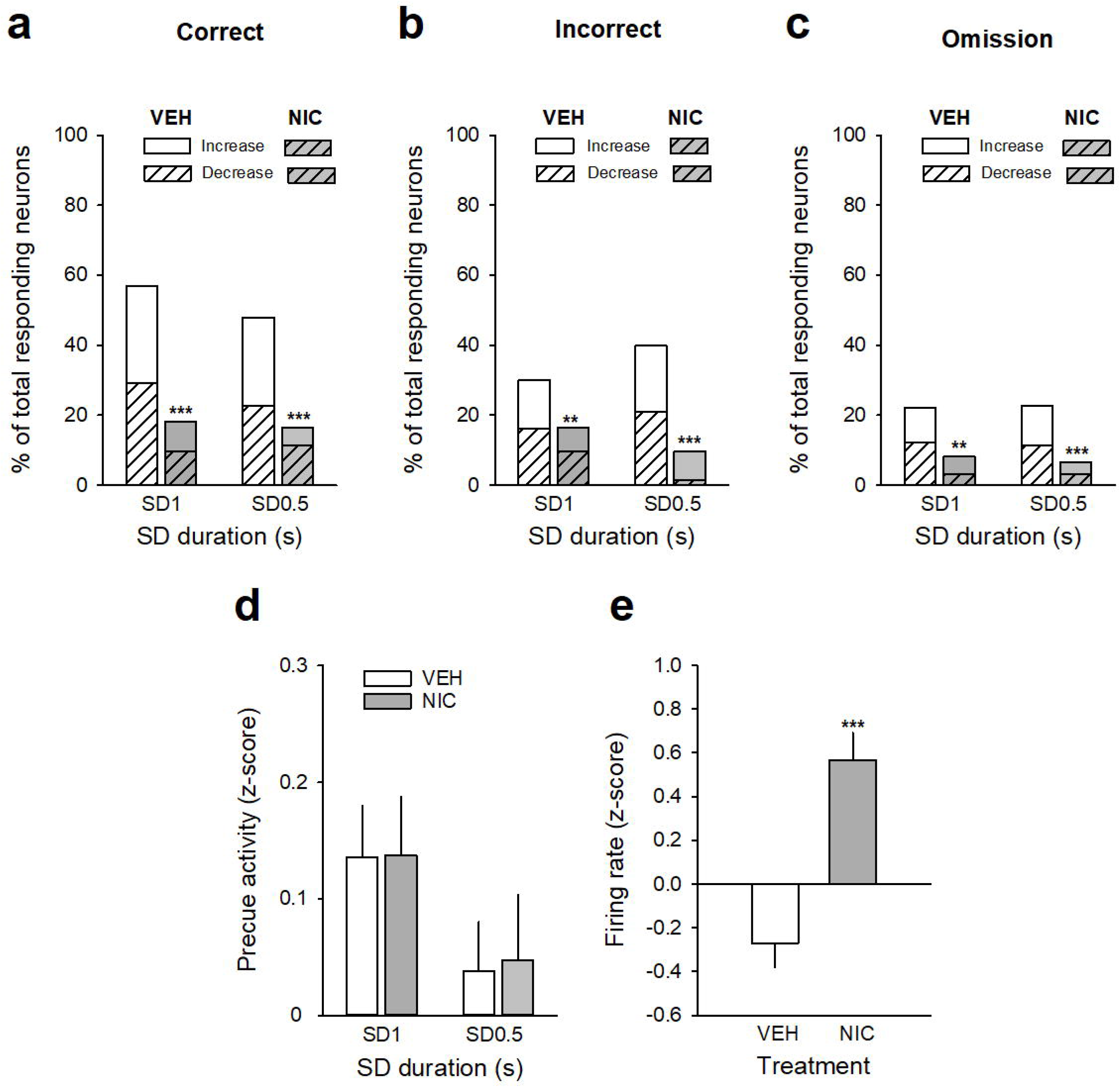
Effects of acute nicotine on vmPFC precue phasic and tonic activities. (**a, b, c**) Percentage of total neurons responding before the cue on correct (**a**), incorrect (**b**), and omission trials (**c**) after a saline injection (VEH, *white bars*) and after a nicotine challenge session (NIC, 10 µg/injection, i.v. 1 min before the session, *grey bars*) during the SD1s and SD 0.5s cue presentation. Excitatory units are shown as solid bars, and inhibitory units are shown as hashed bars. \*\*\**p* < 0.001, different from BL. (**d**) Mean magnitude (±SEM) of precue activity (z-score) of all recorded neurons for all trials combined during SD1s and SD0.5s. (**e**) Mean (±SEM) normalized firing of all recorded neurons (z-scores) during the SD variable session after a saline injection (VEH, *white bars*) and after a nicotine challenge session (NIC, 10 µg/injection, i.v. 1 min before the session, *grey bars*). \*\*\**p* < 0.001, different from VEH.

**Figure 7.**
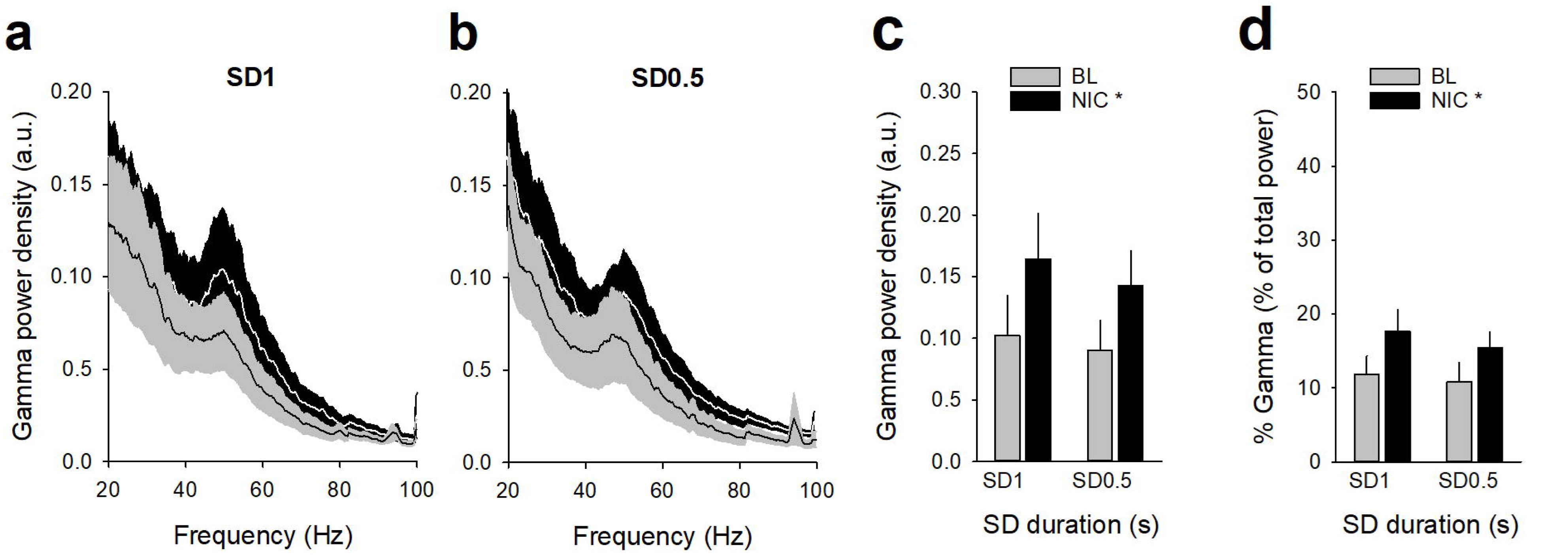
Effects of nicotine on vmPFC gamma oscillations during the SD1s and SD0.5s cue presentation. (**a, b**) Mean (±SEM) LFP power spectrum density across animals after a saline injection (BL, *grey lines*) and after a nicotine challenge session (NIC, 10 µg/injection, i.v. 1 min before the session, *black lines*) during the SD1s (**a**) and SD0.5s (**b**). (**c**, **d**) Mean (±SEM) gamma power density (**c**) and percentage of gamma oscillations (**d**) after a saline injection (BL, *grey bars*) and after a nicotine challenge session (NIC, 10 µg/injection, i.v. 1 min before the session, *black bars*) during the SD1s and SD0.5s. \**p* < 0.05, different from BL.

**Table 4:**
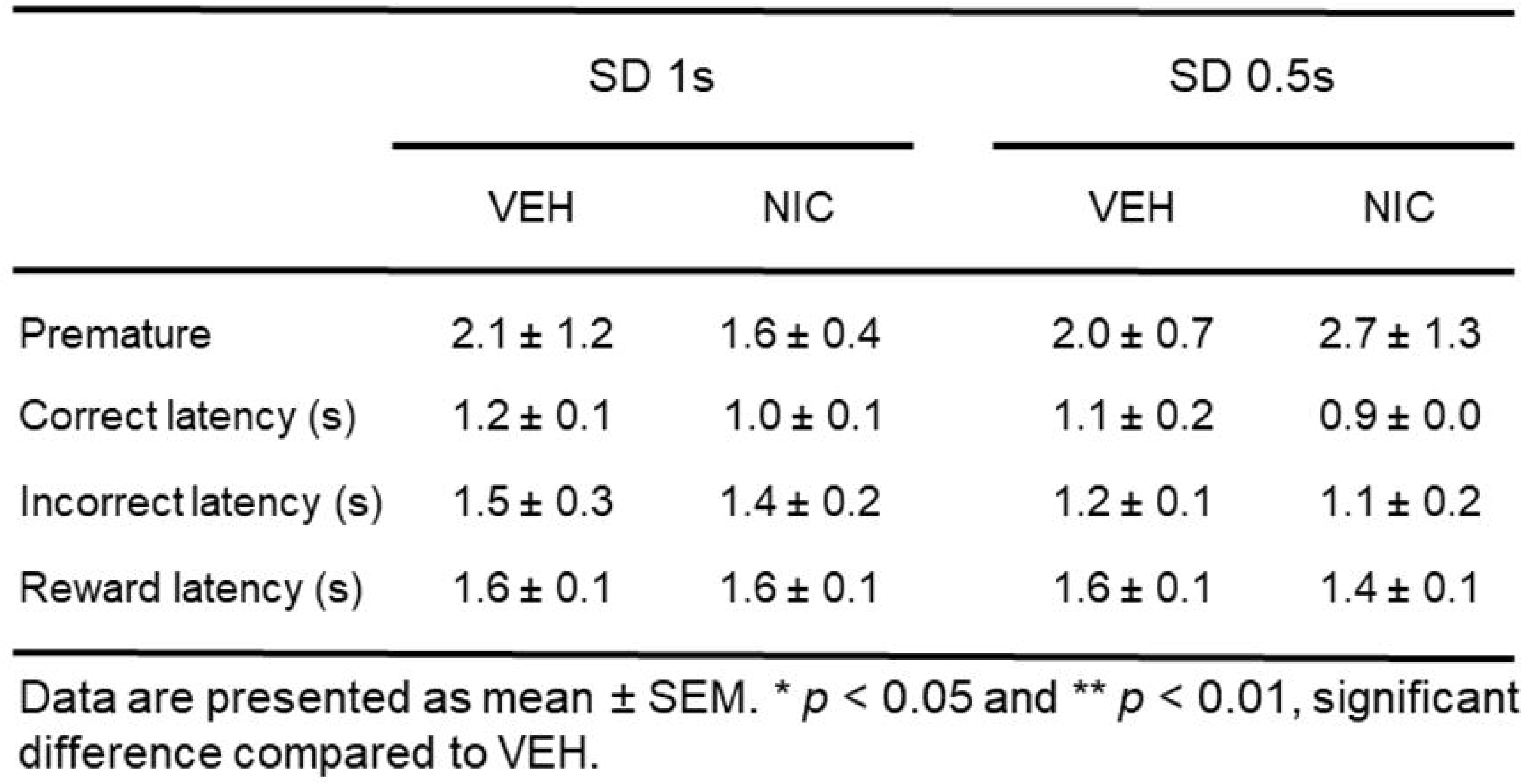
Effects of acute nicotine (1O µg/kg, iv) on 5-CSRTT performances at varying stimulus durations

|  | SD 1s |  | SD 0.5s |  |
| --- | --- | --- | --- | --- |
|  | VEH | NIC | VEH | NIC |
| Premature | 2.1 ± 1.2 | 1.6 ± 0.4 | 2.0 ± 0.7 | 2.7 ± 1.3 |
| Correct latency (s) | 1.2 ± 0.1 | 1.0 ± 0.1 | 1.1 ± 0.2 | 0.9 ± 0.0 |
| Incorrect latency (s) | 1.5 ± 0.3 | 1.4 ± 0.2 | 1.2 ± 0.1 | 1.1 ± 0.2 |
| Reward latency (s) | 1.6 ± 0.1 | 1.6 ± 0.1 | 1.6 ± 0.1 | 1.4 ± 0.1 |

Data are presented as mean ± SEM. \* $p < 0.05$ and \*\* $p < 0.01$ , significant difference compared to VEH.

**Table 5:** Effects of acute nicotine (1O µg/kg, iv) on the number and percentage of vmPFC neurons responding during the precue period

|  |  | Correct trials |  | Incorrect trials |  | Omission trials |  |
| --- | --- | --- | --- | --- | --- | --- | --- |
|  |  | VEH | NIC | VEH | NIC | VEH | NIC |
| SD 1s | Total | 70 (57%) | 22 (18%)<sup>+++</sup> | 37 (30%)<sup>***</sup> | 20 (16%)<sup>++</sup> | 27 (22%)<sup>***</sup> | 10 (8%)<sup>+++</sup> |
|  | Increase | 34 (28%) | 10 (8%)<sup>+++</sup> | 17 (14%)<sup>**</sup> | 8 (7%)<sup>+</sup> | 12 (10%)<sup>***</sup> | 6 (5%)<sup>**</sup> |
|  | Decrease | 36 (29%) | 12 (10%)<sup>+++</sup> | 20 (16%)<sup>**</sup> | 12 (10%) | 15 (12%)<sup>***</sup> | 4 (3%)<sup>++</sup> |
| SD 0.5s | Total | 59 (48%) | 20 (16%)<sup>+++</sup> | 49 (40%) | 12 (10%)<sup>+++</sup> | 28 (23%)<sup>***</sup> | 8 (7%)<sup>+++</sup> |
|  | Increase | 31 (25%) | 6 (5%)<sup>+++</sup> | 23 (19%) | 10 (8%)<sup>++</sup> | 14 (11%)<sup>**</sup> | 4 (3%)<sup>++</sup> |
|  | Decrease | 28 (23%) | 14 (12%)<sup>++</sup> | 26 (21%) | 2 (2%)<sup>+++</sup> | 14 (11%)<sup>**</sup> | 4 (3%)<sup>++</sup> |

\*\*\* $p < 0.001$ and \*\* $p < 0.01$ , significant difference compared to correct trials. ++ $p < 0.01$ and +++ $p < 0.001$ , significant difference compared to VEH.

## DISCUSSION

In this study, we tested whether vmPFC neuronal activity is sensitive to changes in attentional 5-CSRTT performances task using two manipulations: a task manipulation increasing task demand by reducing the cue stimulus duration from 1s to 0.5s, and a pharmacological manipulation administrating an acute nicotine injection (10 µg/inj, i.v.) before the session. As expected from previous research (Amitai and Markou, 2011; Hahn et al., 2002; Levin et al., 2006; Mirza and Stolerman, 1998), increasing task demand led to a decrease in accuracy, while in contrast nicotine injection increased the accuracy. We then recorded and compared vmPFC neuronal activity in these two manipulations that induce opposite effect on attentional performance.

Consistent with previous findings (Donnelly et al., 2015; Kim et al., 2016; Totah et al., 2009), we found that within the vmPFC, a larger proportion of pyramidal neurons showed task engagement with phasic changes in firing rate during the precue period when attention is allocated. These precue responses were observed on each trial (i.e., correct, incorrect or omission trials) and consisted of mixed changes in activity with half of the neurons showing an increase and half of the neurons showing a decrease in firing activity. Importantly, both the proportion of responsive neurons and the magnitude of their responses were graded. That is, fewer units with lower firing activity exhibited phasic responses during the precue period on incorrect and omission trials, compared with correct trials. Thus, vmPFC phasic precue responses appear to be related to the degree of preparatory attention and subsequent response. These precue responses were observed for both SD durations although the number of responsive neurons and the magnitude of precue activity were reduced for the SD0.5s along with poorer attentional performances. Moreover, at the population level of all recorded neurons, the neuronal activity during the precue period was globally higher in SD1s than in SD0.5s, suggesting that enhanced phasic vmPFC activity before the cue presentation facilitates accuracy discrimination.

To further test the link between vmPFC precue activity and attentional performances, we performed optogenetic inhibition of vmPFC pyramidal neurons 2 s before the cue presentation, a time window known to be important for stimulus detection (Luchicchi et al., 2016). Importantly, by doing so in both SD1s and SD0.5s sessions, we found that vmPFC opto-inhibition had different effects on attentional behavior depending on the SD duration. That is, transient opto-inhibition of vmPFC pyramidal before the SD1s cue presentation decreased attention, while the same inhibition protocol before the SD0.5s cue presentation increased it. Moreover, these disruptive effects of vmPFC opto-inhibition on attention affected distinct behavioral parameters. Specifically, in SD1s, vmPFC opto-inhibition before the cue presentation decreased response accuracy by increasing the number of incorrect responses, while in SD0.5s, vmPFC opto-inhibition had no effect on accuracy but increased attentional performances by decreasing the percentage of omissions. This decrease in omission presents as an increase in completed trials, indicating a greater engagement of the rats in the task. Such discrepancies in the effects of optogenetic manipulations of the mPFC in attention have been previously observed in separate studies using different SD cue presentations (Kim et al., 2016; Luchicchi et al., 2016). Importantly, as vmPFC precue activity varied depending on the SD duration, these results suggest that vmPFC opto-inhibition altered attention and attentional parameters in an activity-dependant manner.

Consistent with previous work (Hahn et al., 2002; Levin et al., 2006; Mirza and Stolerman, 1998), we found that nicotine enhanced attentional performances. Surprisingly however, this nicotine treatment almost abolished vmPFC phasic precue responses, an effect due to an overall increase in tonic firing rate of all vmPFC recorded neurons after nicotine injection that persisted during the entire 5-CSRTT session, and could thus limit the detection of phasic changes in firing (Kravitz and Peoples, 2008; Peoples et al., 2007). Consistent with this, others studies have shown that systemic nicotine increased the tonic firing rate of mPFC pyramidal neurons in rats (Bueno-Junior et al., 2017; Zhang et al., 2012), and the activation of frontal networks during attention tasks in humans (Lawrence et al., 2002). Changes in tonic activity of neuromodulators such as acetylcholine or dopamine are also known to play a role in attention by modulating the general efficacy of cortical information processing (Ellwood et al., 2017; Hasselmo and Sarter, 2011; Lohani et al., 2019; Parikh and Sarter, 2008), an effect that could be responsible for the nicotine-induced tonic increase in vmPFC neuronal activity observed here. Together, these results support the role of persistent tonic changes in vmPFC firing rate as a complementary mechanism by which nicotine may facilitate and strengthen information processing in the PFC.

In addition, while increasing the attentional load had no effect on vmPFC gamma oscillations, nicotine injection increased gamma oscillations and associated attention performances at both SD durations. Thus, in contrast to phasic precue vmPFC neuronal activity, the pharmacological pro-cognitive effect of nicotine on gamma activity are independent of the task demand. Such a strengthening of mPFC gamma power and attention has been previously observed after the administration of nicotine or cholinergic agonists (Bueno-Junior et al., 2017; Howe et al., 2017; Pafundo et al., 2013). Gamma oscillations reflect network synchrony as determined by the balance between excitatory and inhibitory (e.g., E/I balance) activity within a region (Sohal et al., 2009; Uhlhaas and Singer, 2013). Interestingly, optogenetic increase of excitatory neurons in the mPFC has been shown to increase spontaneous gamma power and the E/I balance (Yizhar et al., 2011), suggesting that increases in gamma power could reflect an overall increased activity in cortical circuits (Sohal and Rubenstein, 2019). This is consistent with the concurrent increase in vmPFC tonic neuronal activity and gamma oscillation observed in the present study. Moreover, systemic administration of NMDAR antagonists that preferentially bind and inhibit GABAergic neural activity and enhances glutamatergic neural activity, also increased gamma power oscillations in rats mPFC (Carli et al., 2006; Homayoun and Moghaddam, 2007; Molina et al., 2014). Though we did not measure GABAergic interneurons, our finding that nicotine-induced increase in vmPFC tonic neuronal activity and gamma power supports the view that this pharmacological effect reflects an overall increased level of vmPFC circuit activity rather than of rhythmic neural activity that is synchronized across neurons (Sohal and Rubenstein, 2019). Future studies will be needed to clarify the role of the GABA interneurons in the neuronal mechanisms underlying the pro-cognitive effects of nicotine on vmPFC gamma power.

The mPFC is connected with many different brain regions and attention processing likely requires signaling in additional areas. Accumulating evidence across species points to a conserved role of the anterior cingulate cortex (ACC) in normal attention processing (Botvinick et al., 2004; Miller and Cohen, 2001). Indeed, neuronal activity in the ACC is increased during a preparatory sustained attentional state in primates and rodents (Johnston et al., 2007; Totah et al., 2013; Totah et al., 2009), and relatively long-lasting chemogenetic inhibition of this area reduced attention-related performance in mice (Koike et al., 2016). It is thus possible that a functional interaction between ACC and vmPFC neurons could be enhanced after nicotine treatment in the present study. Moreover, the mPFC is reciprocally connected with visual areas (Yeterian et al., 2012), and attentional selection appears to be mediated in part by neural synchrony between neurons in the mPFC and visual areas (Bichot et al., 2015; Gregoriou et al., 2009; Gregoriou et al., 2015; Siegel et al., 2015). Thus, another possibility is that the pro-attentional effects of nicotine could be attributed to an increase of top-down visual processing through the enhancement of synchronization in the gamma frequency range between those brain structures.

Together, these results demonstrate a key role of vmPFC neuronal activity in regulating attentional performances when a cue detection is required to produce an adaptive response. Importantly, using two behavioral manipulations (i.e., an increase in the attentional load and a pharmacological intervention), this study reveals the existence of two distinct neuronal mechanisms operating at different timescales and suggests that allocation of attention could be achieved through multiple complementary neuronal mechanisms within the vmPFC.

## MATERIAL AND METHODS

### Subjects

A total of 28 adult male Wistar rats (250-275 g at the beginning of experiments, Charles River, Lyon, France) were used (8 for the *in vivo* electrophysiology experiment, and 20 for the optogenetic experiment). Rats were housed in groups of 2 and were maintained in a light-(reverse light-dark cycle), humidity- (60 ± 20%) and temperature-controlled vivarium (21 ± 2°C), with water available ad libitum. Animals were food restricted throughout the experiment to maintain at least 95% of their free feeding body weight. Food ration (∼ 14-16 g/rat) were given 3 hours after the end of the session. All behavioral testing occurred during the dark phase of the light-dark cycle. Home cages were enriched with a nylon gnawing bone and a cardboard tunnel (Plexx BV, The Netherlands). All experiments were carried out in accordance with institutional and international standards of care and use of laboratory animals [UK Animals (Scientific Procedures) Act, 1986; and associated guidelines; the European Communities Council Directive (2010/63/UE, 22 September 2010) and the French Directives concerning the use of laboratory animals (décret 2013-118, 1 February 2013)]. The animal facility has been approved by the Committee of the Veterinary Services Gironde, agreement number B33-063-922.

### Five-choice serial reaction time task (5-CSRTT)

#### Apparatus

Height identical five-hole nose poke operant chambers (30 x 40 x 36 cm) housed in sound-insulating and ventilated cubicles were used for 5-CSRTT testing and training (Imétronic, Pessac, France) (Fig. 1a), as previously described (Vouillac-Mendoza et al., 2024). Each chamber was equipped with stainless steel grid floors and a white house light mounted in the center of the roof. Set in the curved wall of each box was an array of five circular holes (2.5 cm sides, 4 cm deep and positioned 2 cm above the grid floor) each equipped with internal light-emitting diodes and an infrared sensor detecting the insertion of the animals’ nose. The opposite wall was not curve and equipped with a delivery port and a drinking cup mounted on the midline. The delivery port was illuminated with a white light diode mounted 8.5 cm above the drinking cup. A lickometer circuit allowed monitoring and recording of licking. Each chamber was also equipped with a syringe pump placed outside, on the top of the cubicle, which delivered sucrose solution into the drinking cup through a silastic tubing (Dow Corning Corporation, Michigan, USA).

#### General training procedure in the 5-CSRTT

During habituation, rats (n = 28) were exposed 60 min to the boxes for two sessions, with the white nose on. During the next three sessions, animals were trained to associate the delivery port with reward without any response requirement. Sucrose rewards (0.1 ml of 10% sucrose) were delivered into the drinking cup with a variable interval of 15 seconds and were signaled by the illumination of the drinking port. Each session terminated after a maximum of 100 sucrose rewards or 60 min, whichever came first. In the next sessions, all five holes were illuminated and rats had to nose pokes into one of them to obtain a sucrose reward as above. Then, only one hole was illuminated and animals were trained to nose poke into that particular hole to obtain a sucrose reward as above.

In the next 5-CSRTT sessions, rats were trained with a maximum number of 100 consecutive self-paced trials per session or a maximum duration of 60 min whichever came first (Fig 1b). The opportunity to self-initiate a trial was signaled by turning on the white light diode in the delivery port. To initiate a trial, rats had to turn off the illuminated port by visiting it (i.e., exit after entering) within one minute. If no visit occurred within the imparted time, the port was turned off automatically and this marked trial onset. After a fixed preparation time of 15 sec, a brief light stimulus was presented behind one of the 5 holes on the opposite curved wall (pseudorandom selection across trials). To receive a reward (0.1 ml of 10% sucrose), animals had to nose-poke the illuminated hole either during the stimulus presentation or within a post-stimulus limited hold period of 5 seconds (Fig. 1b). Correct nose-poke responses immediately turned back on the light in the delivery port and triggered the delivery of sucrose into the drinking cup. Incorrect nose-poke responses in one of the dark holes were not rewarded by sucrose, but were punished by a time-out penalty of 5 seconds signaled by turning on the house light. If animals failed to respond in any of the holes during a trial, this was considered an omission response. Omissions, like incorrect responses, were punished by a signaled 5-s time-out penalty. At the end of the time-out penalty, the house light was switched off and the light in the delivery port was turned back on for the next trial.

During training, the duration of light stimulus was initially set to 30 s and progressively decreased across sessions to 1 s until the subject met performance criteria (omissions < 30%; accuracy > 60%; number of self-initiated trials > 50) or after 10 sessions.

#### Manipulation of attentional task demand

We first investigated the effects of increased attentional task demand by shortening the stimulus duration (i.e., SD) from 1 to 0.5 within the same session. Each SD was tested during 50 trials per session. Animals were tested in this SD variable procedure during several daily sessions until stabilization of performance. Each session terminated after a maximum of 100 sucrose rewards or 60 min, whichever came first.

#### Non-contingent administration of nicotine

Finally, we assessed the effect of a non-contingent administration of nicotine (10 µg/injection, i.v.) 1 minute before the SD variable session. This was done in one session using a SD variable session procedure identical to that described above during a total of 100 trials, except that an intravenous saline injection was administered before the first 50 trials, and a nicotine (10 µg/injection, i.v.) before the last 50 trials. Attentional performance was then measured under baseline condition after a saline injection (VEH) and after a nicotine injection (NIC).

### *In vivo* electrophysiological recordings

#### Surgeries

Animals were anesthetized with a mixture of ketamine (100 mg/kg, i.p., Bayer Pharma, Lyon, France) and xylazine (15 mg/kg, i.p., Merial, Lyon, France) and surgically prepared with an indwelling silastic catheter in the right jugular vein (Dow Corning Corporation, Michigan, USA). Then, under isofluorane anesthesia, an array of 16 teflon-coated stainless steel microwires (2 rows of 8 wires separated from each other by 0.25 mm; MicroProbes Inc, Gaithersburg, MD) were implanted unilaterally in the vmPFC [AP: + 2.5 to + 4.5 mm, ML: 0.3 to 1.2 mm, and DV: - 4.5 mm relative to skull level], as previously described (Abarkan et al., 2023; Girardeau et al., 2019). A stainless-steel ground wire was also implanted 4 mm into the ipsilateral side of the brain, 5 mm caudal to bregma. After surgery, catheters were flushed daily with 0.2 ml of a sterile antibiotic solution containing heparinized saline (280 IU / ml) and ampicilline (Panpharma, Fougères, France). Behavioral testing began 10-14 days after surgery.

#### Neuronal recordings and single unit isolation

Voltage signals from each microwire were recorded, amplified up to 40000X, processed, and digitally captured using commercial hardware and software (OmniPlex, Plexon, Inc, Dallas, TX). Spiking activity was digitized at 40 kHz, bandpass filtered from 250 Hz to 8kHz and captured using commercial hardware and software (Plexon). Behavioral events in the operant task were streamed to the Plexon via TTL pulses delivered from the Imetronic system (Imetronic, Pessac, France) to allow the neuronal data to be accurately synchronized and aligned to these behavioral events. Single units spike sorting was performed off-line: principal component scores were calculated and plotted in a three-dimensional principal component space and clusters containing similar valid waveforms were defined (Offline Sorter, Plexon). The quality of recorded units was ensured with an interspike interval criterion (>1 ms) and a signal:noise criterion (>3X noise band). Neurons were classified into putative pyramidal and interneurons according to the waveform spike width and average firing rate, as previously described (Girardeau et al., 2019; Guillem et al., 2010). Due to the small number of interneurons (*n* = 13), data analysis was exclusively focused on putative the and consisted of a total 184 pyramidal neurons (123 during baseline sessions and 61 during nicotine challenge test session). Electrophysiological data were analyzed using NeuroExplorer (Plexon Inc., Dallas, TX). Each animal was recorded during two sessions: 1) after stabilization of the performance under a SD variable procedure with increased task demand, and 2) after a non-contingent nicotine administration (10 µg/inj, i.v.) 1 min before the session).

#### Neuronal data analysis

Perievent firing rate histograms (PETHs) were used to analyse neuronal responses during the attentional precue period (−2 to 0 s before cue onset) and compared between the three trials types determined by the subsequent cue hole nose poke response (correct or incorrect) or lack of nose poke response (omission) after the cue onset. To examine population activity, PETHs for each unit were normalized in z-score and averaged across different trials (correct, incorrect, and omission), as previously described (Kim et al., 2016). To provide a better visualization of the data, population activity graphs were smoothed with a Gaussian filter.

Neurons were tested for phasic changes during the attentional process by comparing firing rates during the precue period (−2 to 0 s before cue onset) to baseline firing rates during -6 to -4 s before cue onset using a Wilcoxon test (*p* < 0.05), as previously described (Totah et al., 2009). The percentage of neurons was determined for each trials type and each SD duration and compared using the two-proportion z-test. To test for the acute pharmacological effects of nicotine on overall neuronal activity, we compared firing rate (z score) of all recorded neurons during the first 25 min of the 5-CSRTT session after a saline injection to firing during the first 25 min 5-CSRTT session after a nicotine challenge injection (10 µg/inj, i.v. 1 min before the session), using a one-way repeated ANOVA. At the end of the study, histological procedures were used to identify the location of all wire tips used to record neurons (Fig. 2a).

### *In vivo* optogenetic manipulation

#### Surgery and procedures

The effect of vmPFC pyramidal neurons optogenetic inhibition was tested in another group of 20 rats. Under isoflurane anesthesia, light-gated opsins viruses encoding the archaerhodopsin-3.0 protein (*n* = 12; CaMKII-α-eArchT3.0-eYFP; 5.5 x 10^12^ viral molecules/mL; UNC Vector Core) or a control virus (*n* = 8; CaMKII-eYFP 5.5 x 1012 viral molecules/mL; UNC Vector Core) were infected bilaterally in the vmPFC (AP: +3 mm; ML: ± 1.4 mm; DV: -4.5 from skull, angle 10°) to selectively infect pyramidal neurons (Abarkan et al., 2023). A total of 0.5µl/hemisphere was injected at a rate of 0.1 µl/min for 10 min using an Hamilton syringe mounted in an infusion pump after which the injector will be left in place for an additional 10 min to allow virus diffusion. Optic fibers (200/230 µm core diameter Plexon Inc., Dallas, TX) were implanted bilaterally slightly above the injection site (∼0.3 mm) to ensure illumination of the transduced neurons and secured to the skull screws and dental cement. Seven days after surgery, rats were retrained under the SD variable for 20 additional days before being testing for optogenetic manipulations. Opto-inhibition of vmPFC pyramidal neurons (8-11 mW) during 2 s before the cue presentation (Luchicchi et al., 2016) was assessed in two separate SD variable sessions (No light and Light application, random order). Each implanted optic fiber was connected to a LED module (550 nm) mounted on a dual LED commutator connected to an optogenetic controller (PlexBright, Plexon Inc., Dallas, TX).

#### cFos immunohistochemistry

We used cFos immunohistochemistry to check the efficiency of the opto-inhibition of vmPFC neuronal activity. Animals underwent a challenge SD 1s session during which vmPFC pyramidal neurons were opto-inhibited as previously. Ninety min after the start of the challenge session, rats were anesthetized and perfused transcardially with 4% paraformaldehyde (PFA). Brains were removed, serially cut on a cryostat (50 µm) and stored in PBS-0.2% sodium azide as free-floating sections for immunohistochemistry. Brains were first checked for fiber placement and viral expression (Fig. 5b, e). Some floating sections were put in a blocking solution with Normal Donkey Serum 5% in PBS-Triton 0.3% (PBST) during 90 min and then incubated with rabbit anti-GFP antibody (1:2000, Invitrogen, A11122) and mouse anti-CAMKII antibody (1:500, Thermo Fisher, MA1-048) overnight at room temperature (RT). Sections were next washed with PBST and revealed with donkey anti-rabbit antibody conjugated to Alexa 488 (1: 1000, Invitrogen, A21206) and donkey anti-mouse antibody conjugated to A568 (1: 1000, Invitrogen, A10037) for 90 min at RT. Sections were finally mounted in Vectashield medium (Vector Laboratories), coverslipped, imaged on an epifluorescence microscope (Olympus) and analyzed using ImageJ (NIH, USA). Others floating sections were processed for cFos immunohistochemistry analysis as described in details elsewhere (Mihindou et al., 2013; Navailles et al., 2015). Briefly, floating sections were first blocked for unspecific staining in 5%BSA-0.3% triton X100 PBS solution and then incubated over night at RT with rabbit polyclonal anti-cFos antibody (1: 1000; sc-52, Santa-Cruz Biotechnology) diluted in 1%BSA-0.3%Triton-X100 0.01M PBS. On the next day, sections were rinsed in PBS and incubated 30 min in an anti-rabbit polymer system (Anti-rabbit Envision HRP polymer system, Agilent K400311-2). After several rinses, sections were incubated in DAB solution (Envision DAB kit: Agilent K346811-2) to reveal the cFos positive cells. Sections were finally mounted on gelatined coated slides, coverslipped, scanned with a Panoramic Scanner (3D-Histech, Hungary) and analyzed using the Mercator software (Immascope, France). Density of cFos-expressing cells (cells / mm2) was measured by a blind observer in both hemispheres of each rat and averaged.

### Drugs

Nicotine hydrogen tartrate was purchased from Sigma (St. Louis, MO), dissolved in isotonic NaCl (0.9% w/w saline in water), filtered through a syringe filter (0.22 µm) and stored at room temperature. Drug dose was expressed as free base.

### Data Analysis

The following behavioral measures were used for analysis: % accuracy ([100 x correct responses] / [correct responses + incorrect responses]); % omission (100 x number of omissions / total self-initiated trials); premature responses (number of responses that occurred before the presentation of the light stimulus); latency of correct and incorrect responses; and, finally, reward latency (i.e., latency between correct responses and contact with the drinking cup). Behavioral data were subjected to one-way or two-way ANOVAs with SD duration (SD1s or SD0.5s) and nicotine treatment (VEH or NIC) as repeated measures, followed by Tukey post hoc tests where relevant. Electrophysiological study, data were subjected to two-way ANOVAs with trial type (correct, incorrect or omission) as between factor and time as repeated measures, followed by Tukey post hoc tests where relevant.

Percentage of neurons were compared using the two-proportion z-test. Optogenetic study, data were subjected to two-way ANOVAs with virus type (ArChT or YFP) as between factor and light condition (light or no-light) as repeated measures, followed by Tukey post hoc tests where relevant. Statistical analyses were run using Statistica, version 7.1 (Statsoft Inc., Maisons-Alfort, France).

## Author contributions

K.G. designed research and experiments; C.V.M. and K.G. performed *in vivo* electrophysiology and optogenetic experiments, and associated data analysis; N.B. and S.D. performed cFos immunohistochemistry and histological verifications; K.G. wrote the paper.

